# Quantifying social roles in multi-animal videos using subject-aware deep-learning

**DOI:** 10.1101/2024.07.07.602350

**Authors:** Kelly Goss, Lezio S. Bueno-Junior, Katherine Stangis, Théo Ardoin, Hanna Carmon, Jie Zhou, Rohan Satapathy, Isabelle Baker, Carolyn E. Jones-Tinsley, Miranda M. Lim, Brendon O. Watson, Cédric Sueur, Carrie R. Ferrario, Geoffrey G. Murphy, Bing Ye, Yujia Hu

## Abstract

Analyzing social behaviors is critical for many fields, including neuroscience, psychology, and ecology. While computational tools have been developed to analyze videos containing animals engaging in limited social interactions under specific experimental conditions, automated identification of the social roles of freely moving individuals in a multi-animal group remains unresolved. Here we describe a deep-learning-based system – named LabGym2 – for identifying and quantifying social roles in multi-animal groups. This system uses a subject-aware approach: it evaluates the behavioral state of every individual in a group of two or more animals while factoring in its social and environmental surroundings. We demonstrate the performance of subject-aware deep-learning in different species and assays, from partner preference in freely-moving insects to primate social interactions in the field. Our subject-aware deep learning approach provides a controllable, interpretable, and efficient framework to enable new experimental paradigms and systematic evaluation of interactive behavior in individuals identified within a group.

## INTRODUCTION

Automated quantification of social behaviors is increasingly being addressed by software development^1–5^, given the complexity of these behaviors and their importance for fields such as neuroscience, psychology, and ecology^6–8^. However, the automated computational analysis of social behaviors remains challenging. Central to this problem is the identification of roles in a social group with three or more individuals, such as *courting, being courted*, or *idling alone* in a mating choice event (**Fig. 1a-b**). To quantify these social roles, computational approaches should be “subject-aware” – that is, they should assess the behavioral state of every individual one at a time while being cognizant of its social surroundings. In fact, many social behaviors may look the same as non-social behaviors if we do not contextualize one individual in relation to others. For example, in *Drosophila*, one must consider the *courting* male to distinguish a female that is *being courted* from one that is *idling alone* as these two states look similar in terms of female behavior (**Fig. 1b**). Distinguishing social roles like these is key for quantifying mating choice in freely moving partner preference assays. Another challenge arises because every individual can interact with any number of other individuals at any time; thus, the number of all possible interactions increases exponentially with the number of individuals in the group. This causes a combinatorial explosion, which can drastically increase the computational complexity when distinguishing social roles (**Fig. 1c**).

**Figure 1.**
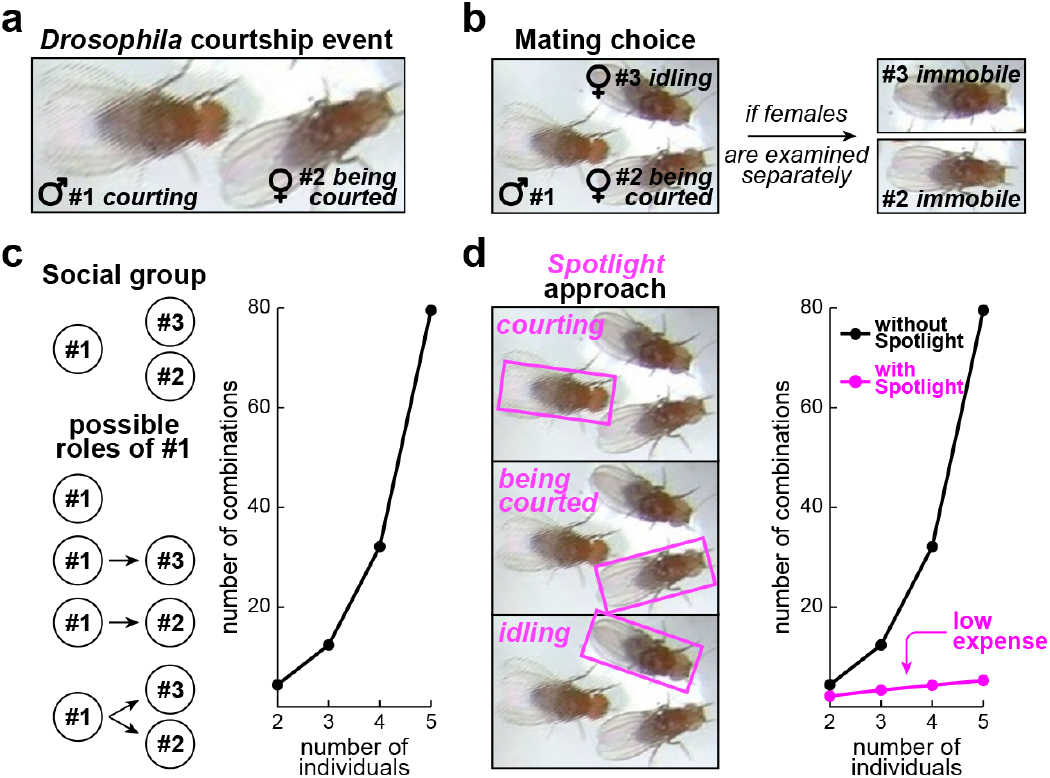
Identifying social roles in *Drosophila* courtship choice assays. **a**, Video snapshot of a courtship event between a male (#1) and a female (#2). **b**, Left: same snapshot, but with wider framing to include a female that is not being courted (#3). Right: the two females appear to behave the same, if they are examined without the social context of the male. **c**, Left: the possible social roles of a given individual, including behaving alone and behaving toward one or two partners. Right: combinatorial explosion with the number of individuals in the video. **d**, Left: using the *Spotlight* approach, each video frame can have different behavioral labels according to the social role of the “main character” in the *Spotlight*. Thus, the social roles of different individuals can be distinguished. Right: the combinatorial explosion is avoided with the *Spotlight* approach.

Several computational tools have been developed to analyze videos of socially interacting animals^9–17^. They include DeepLabCut^12^ and SLEAP^15^, which are pose estimation tools that track user-defined body parts (e.g., head and limbs), also known as “keypoints”. However, tracking multiple animals is just the first step in identifying and then quantifying social behaviors. To that end, other machine learning tools, such as SBeA^11^ and Keypoint-MoSeq^17^, were created to decode keypoint trajectories into motion motifs in social dyads. These approaches are unsupervised: they identify mathematical clusters representing the behavior types (e.g., defined by proximity and relative position of keypoints in space and time), which reduces labor and may identify novel behaviors. However, unsupervised learning tools may erroneously group different behaviors into a single cluster or divide a single behavior into two or more clusters, potentially confounding the interpretations.

Supervised classifications powerfully complement unsupervised classifications, enabling efficient identification of behaviors of interest and thereby capitalizing on decades of neuroscientific experience. This is important since unsupervised methods may fail to classify certain behaviors entirely, rendering certain experiments infeasible with those tools. However, supervised approaches add significant labor to the workflow due to the data-labeling step that is required for the “supervision” or training. Therefore, supervised and unsupervised methods are important complements of each other.

Neither unsupervised approaches nor supervised approaches currently address the combinatorial explosion of computational amount with increasing numbers of subjects in the videos (**Fig. 1c**). Thus, both are limited in identifying social roles (e.g., when the presence of a social behavior relies on directionality between two individuals, such as the fly courtship described above), especially if the number of recorded individuals is increased to three or more. Consequently, current supervised classification tools have only been demonstrated to analyze dyadic interactions or group behaviors in which all individuals perform the same behavior. For example, DeepOF^10^, Simple Behavior Analysis (SimBA)^14^ and Mouse Action Recognition System (MARS)^16^ use body-part keypoint tracking to classify rodent social behavior between two different-looking individuals, typically a dark and a white rodent. Alternatively, DeepEthogram^9^ uses raw pixel data, instead of keypoints, to annotate social proximity events, without however tracking individual identities. This allows DeepEthogram to identify behaviors performed by all animals, but without identifying individual roles or directionality of the interaction. SIPEC^13^ has similar capabilities as DeepEthogram, but it is additionally able to track identities in groups of 2-3 animals, particularly non-human primates behaving in a 3D space. SIPEC can thus classify some group behaviors in non-human primates, such as social grooming, but still without identifying their social roles.

As can be seen from the literature above, an “ecosystem” of software tools is emerging for the study of video-recorded social behaviors, each tool being specialized to a computer vision strategy (e.g., supervised or unsupervised) or experimental setting (e.g., a certain number of animals or specific recording condition). However, we lack a tool that can identify and quantify social roles in groups of three or more individuals while being adaptable to the experimental setting.

We created a subject-aware deep-learning system to meet the challenges above. This system uses an approach called “*Spotlight*” to contextualize every individual in relation to its social surroundings. *Spotlight* guides the deep-learning network toward one individual at a time. As a result, the computational workload increases only linearly with the number of individuals, avoiding the combinatorial explosion problem (**Fig. 1d**). Another way to understand this subject-aware approach is that the individual in the *Spotlight* is the “main character” while its social group is the “stage”. By switching the *Spotlight* one individual at a time, both the “main characters” and the “stage” change accordingly, affecting the behavioral classification. Of note, each main character has a role determined by its relation to the entire remaining group, rather than to each individual other animal – which prevents the combinatorial explosion.

There are two prerequisites for this subject-aware approach. First, the identity of every individual must be accurately tracked over time. Therefore, false detections must be avoided as much as possible, even in situations of close body contact (e.g., social grooming or huddling), or background instability (e.g., variations in lighting or camera angle between or within videos). We achieved reliable identity tracking by customizing a preparatory module called “*Detector*”, which is based on Detectron2, an instance-segmentation package developed by Meta’s Artificial Intelligence team^18^. Second, as the individuals of the social “stage” move around and/ or change their postures, a succession of collective movement patterns occurs, constantly affecting the “main character” and its social role. To classify these movement patterns, we utilized two approaches that we published previously for analyzing single-animal behaviors^19^, called “*Animation*” and “*Pattern Image*”, which feed the deep learning model with comprehensive spatiotemporal information on appearance, postural and movement patterns.

Here we integrated accurate identity tracking (*Detector*), comprehensive information on behavior (*Animation* and *Pattern Image*) and subject-aware deep learning (*Spotlight*), resulting in a novel open-source software called LabGym2 (https://github.com/umyelab/LabGym). LabGym2 addresses the major challenge of social role identification in multi-individual interactions, which had not yet been addressed by the existing tools. Furthermore, LabGym2 is flexible in terms of video angle (e.g., overhead, frontal), background (e.g., bedded cage, petri dish, unstable illumination, or varying landscape), number of animals (e.g., two or more), similarity between animals (e.g., different- or similar-coat color), species (e.g., invertebrates or vertebrates) and experimental paradigms (e.g., partner preference, dyads, social recognition, or interaction with environment features). All of these are accomplished at low manual and computational effort, in addition to avoiding steps that are inherent to unsupervised clustering, particularly the need to simplify metrics for information embedding and dimensionality reduction. Below we report the performance of LabGym2 on videos with *Drosophila*, prairie voles, rats, and mice in the laboratory, as well as macaques in the field. As demonstrated below, LabGym2 may enable the discovery of nuanced behavioral features and potentially influence experiment design in future studies.

## RESULTS

### Avoiding identity switch and tolerating background instability

A prerequisite for distinguishing social roles is to consistently track the individuals over time, avoiding false detections and identity switches. We addressed that challenge by developing a preparatory module called “*Detector*”, which is based on Detectron2, a deep-learning package that detects user-defined objects or animals and generates masks for them^18^. Detectron2 implements a pretrained mask-regional convolutional neural network (mask-RCNN)^20^ and is mostly resistant to background noise (e.g., unstable illumination). However, in our tests, Detectron2 still struggles with mask fragmentation, in which multiple masks erroneously track the same individual, during prolonged periods of social proximity. To address this issue, we incorporated non-maximum suppression and area-based filtering methods to process Detectron2’s masks, followed by appearance- and distance-based tracking algorithms to reinforce the consistency of instance tracking (see Methods). Altogether, Detectron2 and these custom processing steps comprise the *Detector* module.

In addition to the mask fragmentation issue, another source of poor identity tracking is inaccurate or missed detections when the individuals engage in body contact. One solution is to increase the amount of annotated training images, which usually improves detection accuracy in our tests. However, this may defeat the purpose of reducing human labor if the amount of training images is increased too much. To provide users with an idea of the number of annotated frames needed for training a LabGym2 *Detector* to avoid false identity tracking, we tested the relationship between the number of annotated frames and the number of identity switches in different scenarios. First, we tested several 10-minute videos (15,000 frames per video, 25 fps) with five individual adult *Drosophila* of similar appearances that frequently collide or occlude each other (**Fig. 2a**). We found that annotating (with Roboflow^21^) around 300 frames (∼2%) for each video was enough to cover most scenarios of collision or occlusion, and was sufficient to train a *Detector* with 0% identity switch (**Fig. 2a** and **Supplementary Video 1**). Next, we tested several 10-minute videos (18,000 frames per video, 30 fps) with two similar-looking prairie voles, whose identities are visually challenging even to human experts, as prairie voles are highly social and frequently huddle and occlude each other (**Extended Data Fig. 1a**). We again found that annotating around 200 frames (∼1%) for each video eliminated identity switches (**Extended Data Fig. 1b-c and Supplementary Video 2**). Finally, we tested several 10-minute videos (9,000 frames per video, 15 fps) with poor resolution and unstable illumination (**Supplementary Video 3**). Each video contained a rat interacting with a cue light turning on and off prior to reinforcer delivery. Because the rat looked different from the light, we annotated two instance categories (“rat” and “light”) to enable LabGym2’s appearance-based tracking. We found that annotating 100 frames from 5 videos (∼0.2%) was sufficient to train a *Detector* with 0% identity switch. Deploying that same *Detector* to new videos also resulted in 0% identity switch (**Extended Data Fig. 1d**).

**Figure 2.**
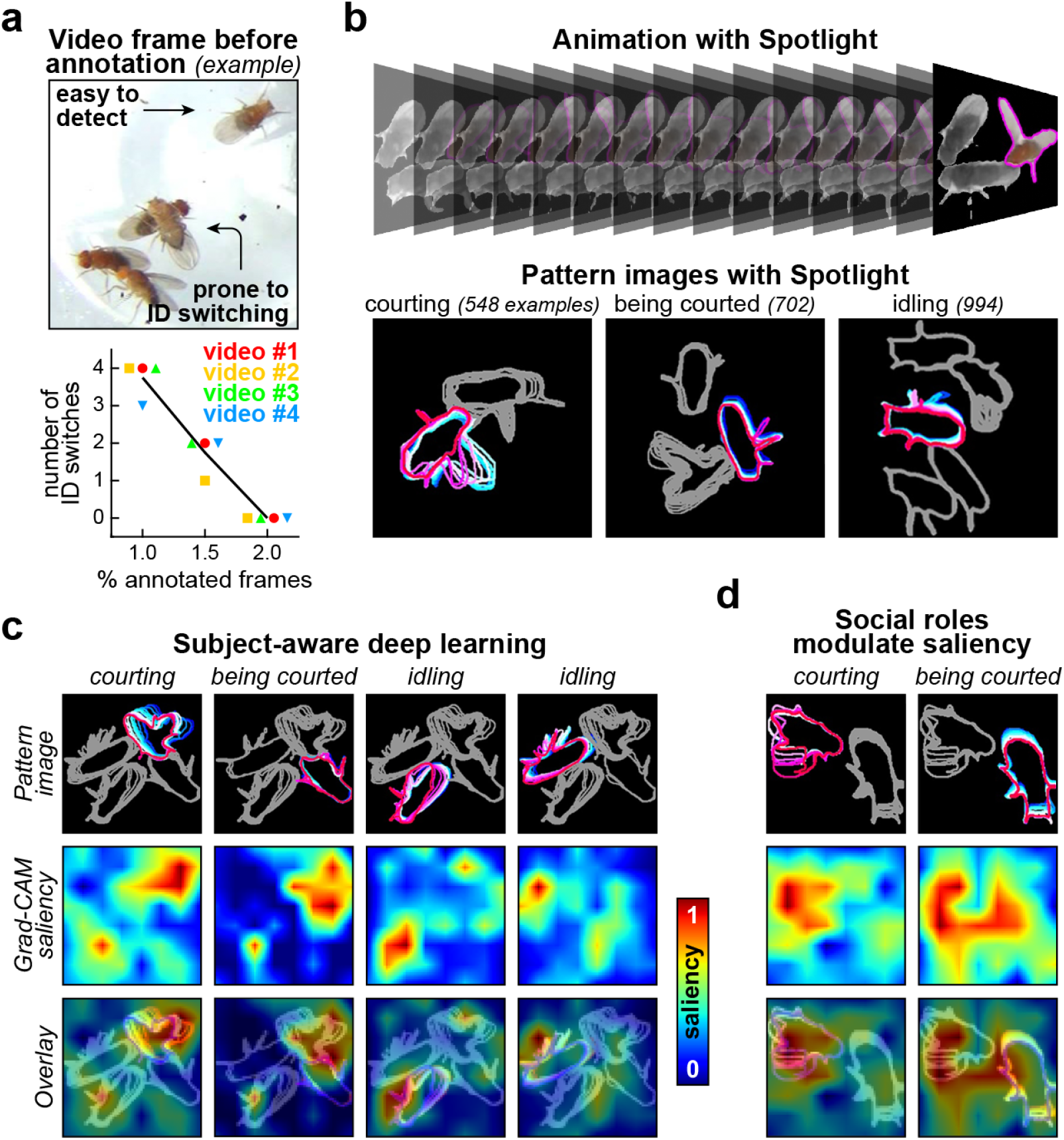
Detecting individuals, extracting motion patterns and using Spotlight to guide deep learning. **a**, Top: video snapshot illustrating visually challenging scenarios of occlusion or collision between individual flies. One can train the *Detector* by manually annotating the individuals in frames like this. Bottom: identity switching is eliminated with a relatively low percentage of manually annotated frames per video. **b**, Examples of *Spotlight Animation* and *Pattern Images* showing fly behaviors, from ∼0.6-s time-windows in the video (15 frames, 25 fps). Indicated are the numbers of behavioral examples used to train the *Categorizer* in this analysis. **c**, Post-hoc measurement of the attention of the *Categorizer* toward regions in the images. *Pattern Images* are overlaid with saliency maps from Gradient-Weighted Class Activation Mapping, or Grad-CAM. Warm colors tend to follow the *Spotlight* individuals, demonstrating subject-aware deep learning. **d**, The same courtship event produces different saliency maps for c*ourting* or *being courted*, demonstrating that social roles can differentially modulate the attention of the neural network.

### *Spotlight*-guided deep learning of social roles

After using *Detector* to track the individuals, the information on their behavioral state is extracted in preparation for subsequent analysis. That information is obtained through two approaches called *Spotlight Animation* and *Pattern Image* (**Fig. 2b**). We recently published the approaches of *Animation* and *Pattern Image* for collecting spatiotemporal information of single behaving animals from user-defined time windows in the video, each of which spans the approximate duration of a behavior episode^19^. *Animation* extracts the pixels occupied by every individual on a per-frame basis, capturing features like body shapes, fur colors and limb positions, as well as kinetic patterns like postural changes. *Pattern Image*, on the other hand, displays the body contours of every individual during the behavior episode. To efficiently reduce the dimension of the time-series data, these contours are merged into a single motion-conveying image, one image per time window. Thus, *Animation* and *Pattern Image* provide a dataset focused on the individuals and their behaviors.

We integrated *Animation* and *Pattern Image* with *Spotlight*: the subject-aware approach that we developed in this study. *Spotlight* adds a color code into *Animation* and *Pattern Image* to differentially mark the individual being analyzed versus its social peers or surroundings (**Fig. 2b**). That color-coding is performed one individual at a time, so that a different individual is colored in the same *Animation* and *Pattern Image* in each iteration. In other words, the same social interaction event (i.e., the same time window in the video) can yield a different behavioral classification depending on which individual is in the *Spotlight*, providing the basis for social-role identification with low computational effort (**Fig. 1c-d**). In addition, users can define the “social distance” threshold (see Methods), so that non-*Spotlight* individuals that are distant from the social interaction event are excluded from *Animation* and *Pattern Image*, further sparing computational effort, and reducing chance of error.

Once the *Spotlight Animation*s and *Pattern Images* are ready, LabGym2 proceeds to behavioral categorization using a deep-learning model previously published by our group, called “*Categorizer*”^19^. For *Categorizer* training, *Spotlight Animation* and *Pattern Image* examples are manually sorted into different behavioral categories, resulting in a neural-network training dataset (**Fig. 2b**). The *Categorizer* is then trained based on that dataset, and later deployed to unseen videos for automated analysis. *Spotlight Animation* and *Pattern Image* can be used in combination or separately to train a *Categorizer*, depending on the need. For example, *Animations* may be dispensable if pixel-based information, such as fur color and body features, do not contribute to social role identification.

We used Gradient-Weighted Class Activation Mapping, or Grad-CAM^22^, to test whether the *Spotlight* approach indeed guides the attention of the *Categorizer* as intended. Grad-CAM produces heatmaps whose warm colors indicate saliency, i.e., important regions in the images being analyzed. First, we sorted two datasets of *Spotlight Pattern Images*, one for training and another for testing, into three behavioral categories in *Drosophila*: *courting, being courted* and *idling* (**Fig. 2b**). We then trained a

*Categorizer* based on the training dataset and used Grad-CAM to analyze the *Categorizer’s* attention toward the testing dataset. This resulted in classification saliency maps (**Fig. 2c**). We found that *Spotlight* indeed guided the *Categorizer’s* attention toward the “main characters”, thus distinguishing their social roles in various situations, including different numbers of individuals. Intriguingly, the salient areas (warm colors) of the “*courting*” and “*idling*” maps corresponded mostly to the individuals performing such behaviors (**Fig. 2c**). In contrast, the salient areas of “*being courted*” overlapped with the two subjects participating in the courtship event (**Fig. 2c-d**). This suggests that *Spotlight* goes beyond just highlighting “main characters”; it guides the attention of the deep-learning model in adaptable manners depending on the behavior, either toward solitary individuals or interacting pairs.

### Quantifying abnormal social choice in a *Drosophila* model of neurodegeneration

Exposing *Drosophila* individuals to multiple potential partners can reveal behavioral repertoires that are not observable in numerically simpler paradigms (i.e., dyads), and provides mechanistic insights into social choice^23^. Distinguishing social roles in such multi-individual interactions allows us to identify which individual is being chosen among all potential partners. We analyzed a *Drosophila* courtship choice assay, in which a male fly was grouped with two younger and two older female flies. Wild-type control males typically spend more time courting younger females, whereas males expressing certain neurodegenerative proteins in the central nervous system lose their preference for younger females^23^. This assay is computationally challenging because it involves five individuals and, therefore, a high number of combinations of possible interactions (80 in total; **Fig. 1c**), which no existing tool can handle.

LabGym2 training comprises two supervised steps: training *Detectors* for animal detection and training *Categorizers* for behavioral classification and quantification. Throughout the studies in this paper, the *Categorizer* used separate datasets for network training (“training videos”) and computerized labeling (“testing videos”). For each video in our multi-individual *Drosophila* assay, we first annotated ∼2% of the frames to cover most of the visually challenging scenarios, and trained a *Detector* on the annotated frames (**Fig. 2a**). We then used the trained *Detectors* to generate *Spotlight Animations* and *Pattern Images* from the training videos (**Fig. 3a**) and sorted them into six categories, including solitary and role-specific social behaviors (**Fig. 3b**). Finally, we trained a *Categorizer* and deployed it to testing videos (**Fig. 3c-d**).

**Figure 3.**
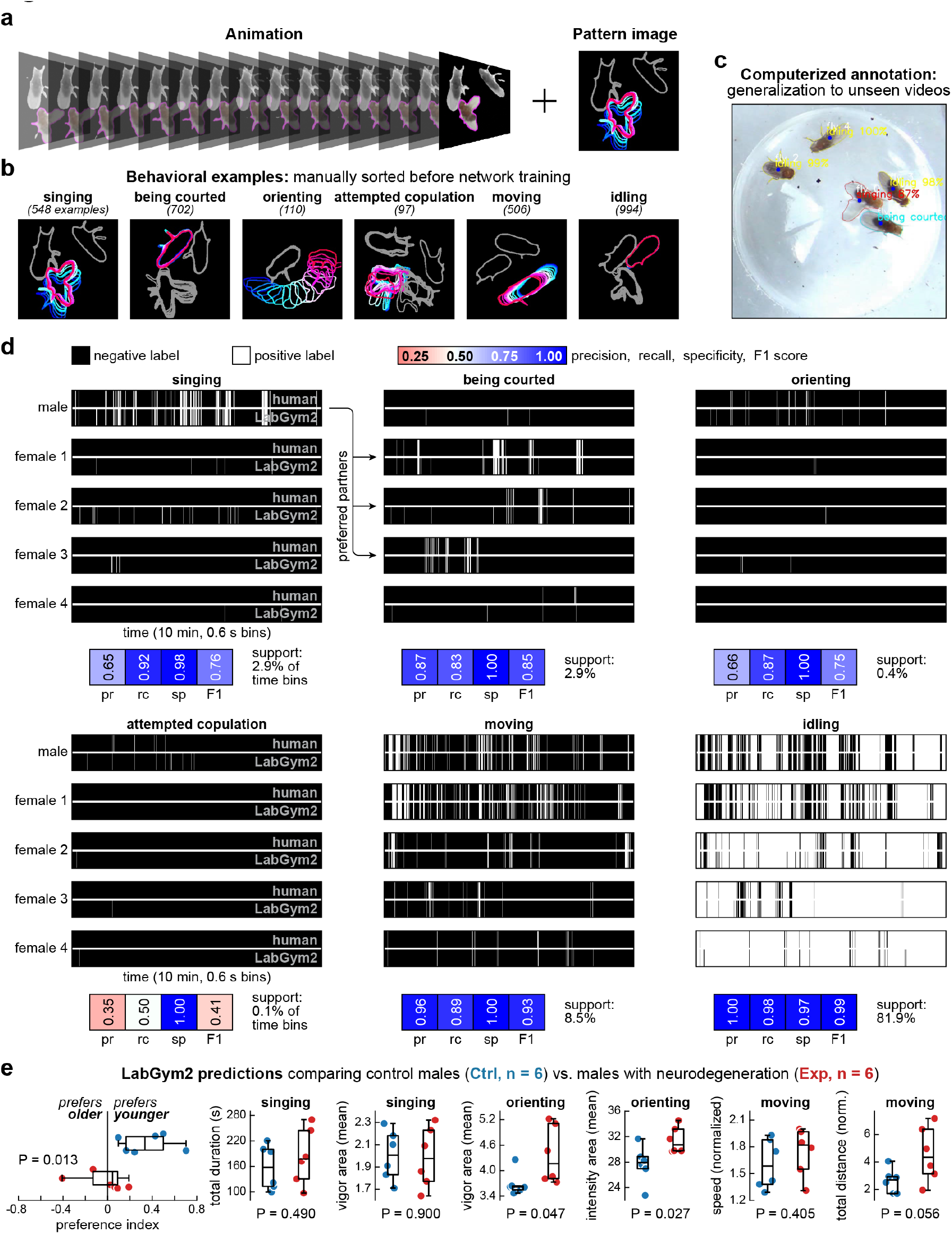
Quantifying abnormal social choice in a *Drosophila* model of neurodegeneration. **a**, *Spotlight Animation* and *Pattern image*. Both contribute to accuracy in this application. These examples are from a ∼0.6-s time-window in the video (15 frames, 25 fps). **b**, Example behaviors in a *Drosophila* social choice assay with one male and four females. Behavioral examples like these are manually sorted for *Categorizer* training. Indicated are the numbers of behavioral examples used for training. The examples may contain less than five individuals because some of them can be excluded based on user-defined “social distance” criteria. **c**, Once trained, the *Categorizer* can generalize the behaviors to unseen videos, as illustrated by this snapshot. **d**, Each subplot shows behavioral annotation timelines from five flies (black-and-white bars) and aggregate evaluation metrics from all animals (red-blue scale). **e**, LabGym2 quantification outputs showing statistical comparisons (unpaired t-test) between wild-type control (Ctrl) and neurodegeneration experimental (Exp) groups in terms of various quantitative measures (see Methods). Each dot represents a male fly, box plots show the inter-quartile range including the median, and whiskers show extreme non-outlier values.

We found that the *Categorizer* we trained for this social assay achieved good accuracy across behaviors (**Fig. 3d**). For example, the human-annotated *singing* behaviors of the male mostly matched the *being courted* behaviors distributed in the females, especially three of them, which could be interpreted as preferred partners (**Fig. 3d**, see arrows, and **Supplementary Videos 4-5**). The *Categorizer* was also able to identify other social roles, like *idling*, as well as non-social behaviors, like *moving* (**Fig. 3d**), confirming its capability to identify both social roles and non-social behaviors in multi-individual interactions.

Finally, we tested the ability of the same *Categorizer* to quantify abnormal social choice. For that, we used 12 testing videos (10 minutes per video) from our previous study^23^, all of them with one male, two younger females and two older females. Six videos contained males with experimental neurodegeneration, and six others contained wild-type controls. The *Categorizer* was sensitive enough to detect abnormal courtship preference in males with neurodegeneration (**Fig. 3e**), consistent with the previous report^23^. Interestingly, the *Categorizer* not only corroborated knowledge, but also revealed new findings. The behavioral deficits in the experimental males were confined to social preference, as we observed no impairments in the duration or vigor of *singing* behaviors (**Fig. 3e**), suggesting that males with neurodegeneration lose social selectively, but maintain their reproductive drive. On the other hand, males with neurodegeneration showed elevated vigor and intensity of *orienting* behaviors (**Fig. 3e**), as well as a statistical trend to increased *moving* distance without changes in speed (**Fig. 3e**), suggesting that males with neurodegeneration show higher locomotion and lower efficiency at targeting the appropriate mates. These quantifications would be impractical via manual scoring and impossible for other computational tools. These results demonstrate LabGym2’s unique capability in quantifying social roles in multi-individual interactions, including derived measures of behavior such as vigor and speed (see Methods).

### Quantifying social changes in prairie voles exposed to early-life sleep disruption

Prairie voles (*Microtus ochrogaster*) are a wild rodent species known for their high incidence of affiliative behaviors and monogamous bonding^24^. Prairie vole social behavior is also measurable in the laboratory, with the aim of studying the neural and hormonal correlates of affiliation^25^. Manual behavioral scoring remains prevalent in prairie vole studies, as their social behaviors are often complex, sequenced, and subtle. Therefore, video recording methods in prairie voles are ripe for integration with deep-learning applications that can “see” these behaviors just as human experts. It has previously been shown that prairie voles exposed to early-life sleep disruption (ELSD) displayed persistent changes in social behaviors into adulthood, including decreased huddling during a partner preference test^26^ and altered body orientation and other social approach behaviors in the first hour of cohabitation with a partner^27^.

Our recent study tracked body-part keypoints in male-female pairs of prairie voles^27^. In that study, the individuals were confined to non-overlapping spaces and filmed from an overhead angle, which yielded continuous variables of body orientation toward the conspecific. Here, we used a separate dataset of overhead videos with fully interactive male-female pairs, i.e., without spatial restrictions (**Fig. 4a-b**; see Methods). We then used LabGym2 to identify social behaviors, instead of collecting posture-related variables for subsequent coding-based analysis like the previous study^27^. We also challenged LabGym2 to predict sex based on social roles, without visual identifiers distinguishing male vs. female. To our knowledge, this is the first report of software that can quantify social roles in prairie vole dyads.

**Figure 4.**
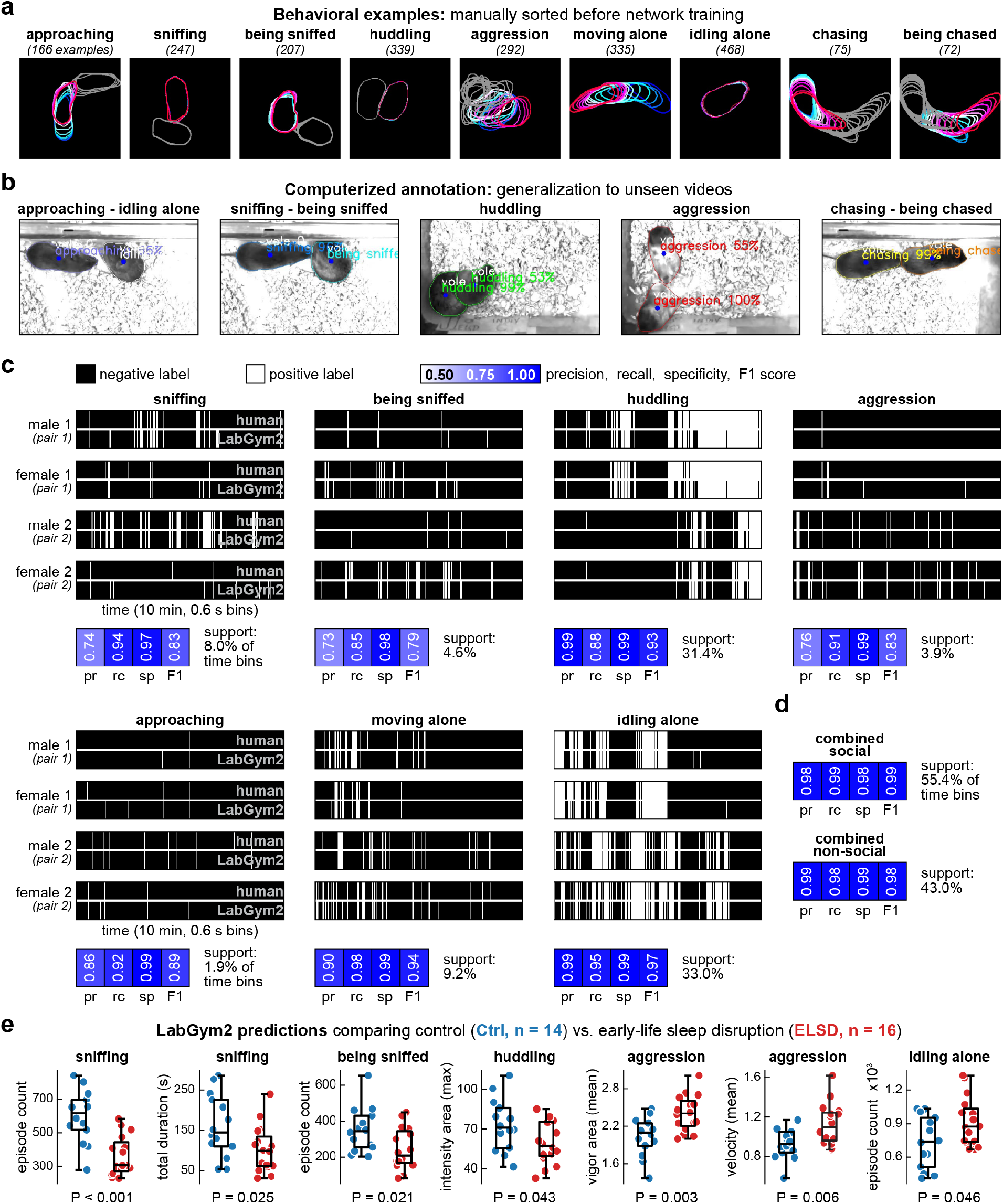
Quantifying social changes in prairie voles exposed to early-life sleep disruption. **a**, Example behaviors in male-female prairie vole pairs. *Spotlight Pattern images* like these are manually sorted for *Categorizer* training. Indicated are the numbers of behavioral examples used for training. *Spotlight Animations* were not necessary in this application as individuals looked very similar, which was not detrimental to accuracy. **b**, Once trained, the *Categorizer* can generalize the behaviors to unseen videos, as illustrated by these snapshots. **c**, Each subplot shows behavioral annotation timelines from two male-female pairs (horizontal black-and-white bars) and aggregate evaluation metrics from all animals (white-blue scale). **d**, Accuracy increases to ∼99% if the behavioral category system is simplified. **e**, LabGym2 quantification outputs showing statistical comparisons (unpaired t-test) between control (Ctrl) and early-life sleep disruption (ELSD) groups in terms of various quantitative measures (see Methods). Each dot represents a prairie vole, without specifying males or females. Box plots show the inter-quartile range including the median, and whiskers show extreme non-outlier values.

We sorted the *Spotlight Patten Images* into nine behavioral categories for the *Categorizer* training (**Fig. 4a-b**). We did not include *Spotlight Animations* because the male-female prairie vole pairs in our videos looked very similar, and adding the pixel-containing *Spotlight Animation* approach did not contribute further to *Categorizer’s* accuracy. We then focused on the seven most frequent behaviors in our dataset and found that the *Categorizer* mostly matched the human labeling (**Fig. 4c**). That accuracy included social role identification within each male-female pair: note the complementarity between *sniffing* in the male and *being sniffed* in the female of the same pair (**Fig. 4c** and **Supplementary Videos 6-7**). Social behaviors without role identification (e.g., *huddling*), and non-social behaviors (e.g., *idling alone*), were also accurately tracked. The computerized labeling accuracy increases to 99% if a simplified behavioral category system is used (**Fig. 4d**). This shows that the user can train the *Categorizer* according to their experimental needs, for example by finding compromises between enriched category systems with infrequent – and visually challenging – behaviors (e.g., compare *approaching* vs. *sniffing* in **Fig. 4b**) and category systems with lower complexity but higher accuracy (**Fig. 4d**).

LabGym2 also detected social changes caused by ELSD (**Fig. 4e**). According to a previous study^26^, ELSD causes lasting alterations in prairie vole sociality, including lower incidence of affiliative behaviors into adulthood. We found that the incidence of *sniffing* and *being sniffed* was indeed significantly lower in ELSD animals compared to controls (**Fig. 4e**). The incidence of *huddling* did not differ between experimental and control groups, but *huddling* in ELSD animals did show lower intensity area than controls (intensity area is another derived behavioral metric calculated by LabGym2; see Methods). In addition, ELSD animals showed higher vigor and velocity in their *aggression* bouts and spent more time *idling alone* than controls, suggesting diminished interest in affiliative social interactions (**Fig. 4e**). Thus, ELSD prairie voles are overall more aggressive and show less affiliative social behaviors. These findings both replicate and expand on previous results that used manual behavioral scoring^26^. This, along with the *Drosophila* assay described above, confirms the diverse applicability of LabGym2: from mating choice studies with three or more individuals to dyad studies with visually challenging behavioral categories.

### Quantifying changes in social recognition in mice fed with high-fat diet

The *Detector* module of LabGym2 performs instance segmentation, which can also detect inanimate objects. This further diversifies the applicability of LabGym2 to animal-object interactions. We tracked behaviors (*climbing, sniffing, rearing*, and *moving*) of an individual mouse relative to either of two fixed-location objects: one with and another without a hand-drawn cross (referred to as “cross-marked” or “blank”, respectively) (**Fig. 5a-d**). These object-referenced behaviors were accurately tracked by the *Categorizer* (**Fig. 5e-f** and **Supplementary Videos 8-9**). It is noteworthy that these objects were identical and only differentiated by the hand-drawn cross. More evident features, such as shapes and colors, should further facilitate instance detection, making *Detector* applicable to videos with movable objects, for example in environmental enrichment studies.

**Figure 5.**
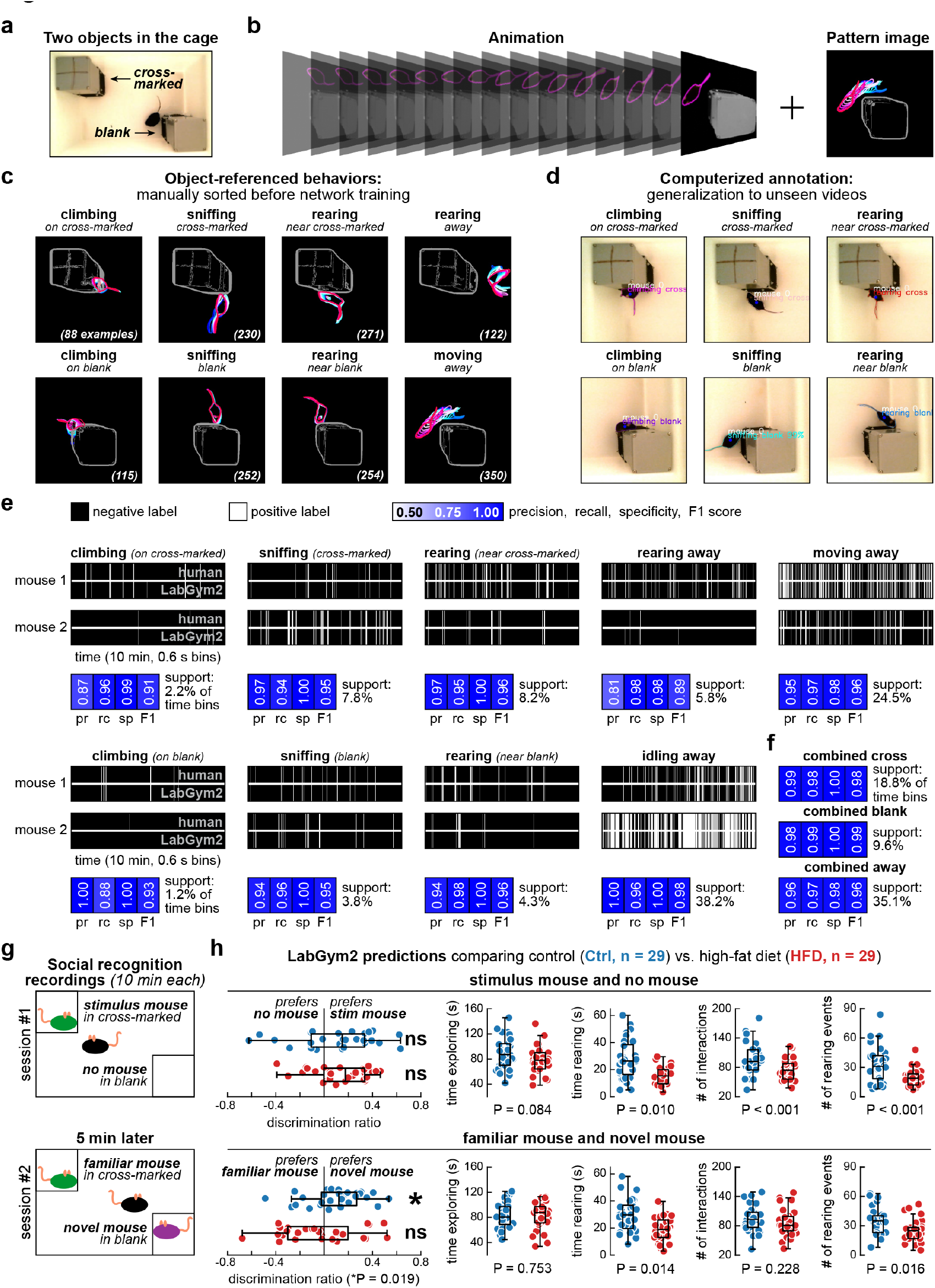
Quantifying changes in social recognition in mice fed with high-fat diet. **a**, Video snapshot showing a test mouse behaving relative to two objects in the cage. **b**, *Spotlight Animation* and *Pattern image*. Both contribute to accuracy in this application. These examples are from a 1-s time-window in the video (15 frames, 15 fps). **c**, Example object-references behaviors, which are manually sorted for *Categorizer* training. Indicated are the numbers of behavioral examples used for training. **d**, Once trained, the *Categorizer* can generalize the behaviors to unseen videos, as illustrated by these snapshots. **e**, Each subplot shows behavioral annotation timelines from two mice from separate recordings (horizontal black-and-white bars) and aggregate evaluation metrics from both animals (white-blue scale). **f**, Accuracy increases to ∼100% if the behavioral category system is simplified. **g**, The objects in this application are mouse containers, making this a social recognition application. Illustrated is the experimental paradigm, which measures social recognition based on the preference for a familiar vs. novel social stimulus. **h**, LabGym2 quantification outputs showing statistical comparisons (unpaired t-test) between control (Ctrl) and high-fat diet (HFD) groups in terms of various quantitative measures (see Methods). Each dot represents a mouse, box plots show the inter-quartile range including the median, and whiskers show extreme non-outlier values.

In the social recognition assay^28,29^, a freely moving “test mouse” interacts with the two objects (**Fig. 5a**). Each object is a mouse container that can either be empty or hold a stimulus mouse. The experiment started with a social stimulation session (session #1, 10 minutes), in which a stimulus mouse was placed in one of the containers and the test mouse was allowed to explore (**Fig. 5g**). That was followed by a memory retention interval (5 minutes) when both animals were removed from the cage. The two initial mice were then put back into the cage for session #2 (10 minutes) along with a novel mouse in the previously empty container (**Fig. 5g**). From this paradigm we obtained the discrimination ratio (**Fig. 5h**), which measures the relative time the test mouse spends near each container (see Methods). By analyzing LabGym2’s outputs, we observed that the control group of test mice preferred the container holding the stimulus mouse in session #1 and shifted the preference to the container holding the novel mouse in session #2. An experimental group of mice fed with high-fat diet (HFD) behaved statistically the same as controls in session #1, suggesting that HFD mice were as socially active as controls. However, in session #2 the HFD animals showed no preference for either container, suggesting impaired social recognition (**Fig. 5h**). This is consistent with previous studies showing that HFD mice have reduced preference for novel social stimuli^30,31^, which in turn indicates deficits in social recognition memory^32^.

Although the discrimination ratio data could have been obtained with other software tools (e.g.,DeepLabCut^12^ or SLEAP^15^), the novelty here is that we quantified social recognition in more detail, thanks to LabGym2’s ability of annotating the object-referenced postural motifs of *climbing, rearing* and *sniffing*. In fact, HFD and controls did not differ in terms of general object exploration but did differ in terms of these postural motifs (**Fig. 5h**). For example, HFD mice spent less time rearing, which replicates previous studies on HFD rearing behavior on the open field and elevated plus maze^33–35^. Thus, in addition to acquiring discrimination ratio data, LabGym2 provided more detailed behavioral analysis, which would not have been accomplished with other tools.

### Distinguishing social roles of Japanese macaques in the field

The social behaviors of animals in the wild pose a significant challenge for computational analysis. Unlike controlled laboratory settings, the wild environment presents a multitude of variables, ranging from environmental factors to unpredictable behavioral dynamics^36–38^. Moreover, the inconsistency in recording locations and the low chances of encountering the wild animals can make the videos highly variable and low in quantity. To assess the applicability of LabGym2 in analyzing social behaviors of wild animals, we analyzed eight videos (∼30 seconds for each) of Japanese macaques in their natural habitat^39,40^, which were divided into five training videos and three testing ones. The shooting angle varied greatly across these videos, and we challenged LabGym2’s *Categorizer* by intentionally including testing videos that are very different from the training videos (**Fig. 6a**).

**Figure 6.**
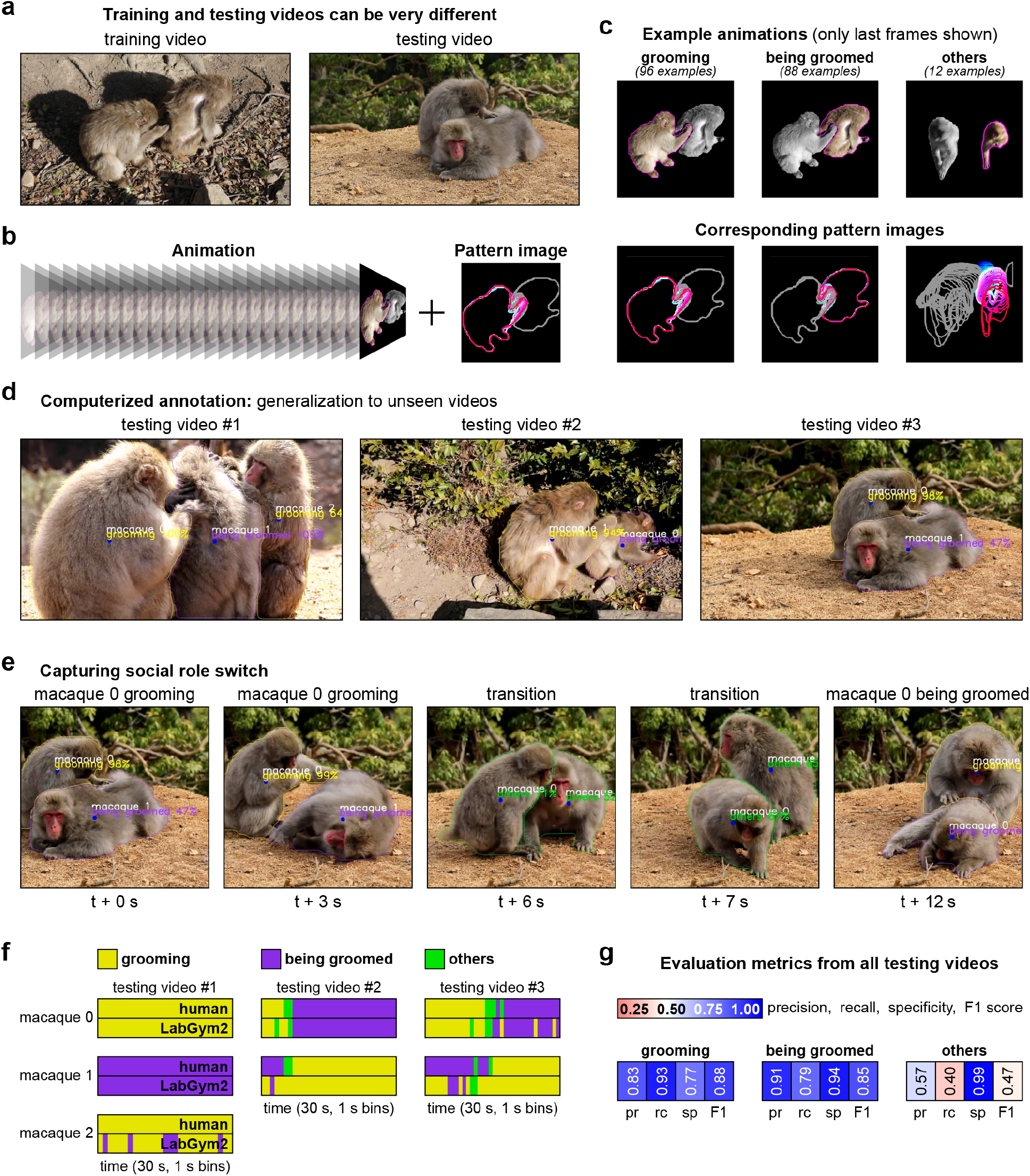
Distinguishing social roles of Japanese macaques in the field. **a**, Two video snapshots from a training and a testing video, with very different shooting angles (fields of view) and background environments. **b**, *Spotlight Animation* and *Pattern image*. Both contribute to accuracy in this application. These examples are from a 1-s time-window in the video (25 frames, 25 fps). **c**, Example macaque behaviors, which are manually sorted for *Categorizer* training. Indicated are the numbers of behavioral examples used for training. **d**, Once trained, the *Categorizer* can generalize the behaviors to unseen videos with different fields of view and backgrounds. **e**, LabGym2 can capture shifts between social roles, as illustrated by this temporal sequence of snapshots. **f**, Each subplot shows behavioral annotation timelines from seven macaques from three separate videos (different colors represent different behaviors). The videos correspond to the snapshots in panel **d.** **g**, Aggregate evaluation metrics from all three videos, parsed by behavior (red-blue scale).

Social cohesion in Japanese macaques is primarily maintained through grooming behavior, especially among females and juveniles^41,42^. Therefore, in this analysis we focused on three categories of macaque behaviors: the social roles of *grooming* and *being groomed*, and a category of unspecified behaviors called “*others*” (**Fig. 6b-c**). We found that, with as few as 12∼96 training examples per behavioral category, the *Categorizer* achieved over 80% accuracy in distinguishing macaque social roles in very different testing videos (**Fig. 6d-f**; **Supplementary Videos 10-11**). These results demonstrate LabGym2’s capacity in quantifying social roles of animals in the field, despite the variability of shooting angles and limited quantity of videos.

## DISCUSSION

In this study, we present LabGym2, a subject-aware deep-learning system for quantifying social roles in multi-animal interactions. It builds on the open-source tool LabGym1^19^, which we previously developed for automated markerless behavioral analysis across species. LabGym1 can only analyze behaviors of single animals on a static background, which precludes studies in natural environments and social interactions. LabGym2 resolves these limitations: it faithfully tracks animals in dynamic backgrounds and uses a *Spotlight*-guided analysis framework to determine the context-specific behavioral role for every individual in a social group, one at a time. LabGym2 enables the automation of challenging analyses that other existing tools are unable to perform, which may lead to the creation of new behavioral paradigms and enable the discovery of new behaviors.

LabGym2 is applicable to a broad range of social and object exploration studies in diverse species and various recording conditions, partly because it tracks and segments individual animals or objects without restrictions on animal species or object features. Moreover, LabGym2 is tolerant to background or lighting variations, expanding its applicability to natural environments, such as social interactions in the wild. Importantly, behavioral identification in LabGym2 does not rely on body postures. Instead, *Spotlight Animations* and *Pattern Images* feed the deep neural network with complete information of each behavioral event, such as body colors and shapes, limb positions, and movement patterns. Such holistic assessment makes it possible for LabGym2 to identify posture-irrelevant behaviors (e.g., color change in chameleons), which are challenging for existing tools that rely on posture information for behavioral identification. In addition to identifying behaviors, LabGym2 captures the kinetic and dynamic features of these behaviors into quantitative measures, such as intensity, acceleration, speed, and vigor. Thus, LabGym2 can quantify behaviors in a nuanced manner, rather than just timestamping these behaviors.

The *Spotlight* approach is a novel deep-learning approach with an efficient, controllable, and interpretable framework. Deep-learning models, despite being powerful, are known to be “black boxes” that are difficult to control or interpret^43^. The *Spotlight* approach addresses this problem by effectively providing a control mechanism. By allowing the dynamic switch between the “main character” and its “stage”, *Spotlight* guides the deep-learning network to focus on key information from the video frames, enabling the network to recognize the roles of each detected individual. These roles can be social roles, which is the focus of this study, but can also include other kinds of “roles”, such as neural activity patterns in calcium imaging. Therefore, the *Spotlight* approach can also be applicable to other computational problems more generally, including in imaging studies involving multiple objects and roles. Finally, the *Spotlight* approach can also be viewed as an efficient decomposition mechanism. It converts the complex problem of (any-in-n)-versus-(any-in-n) into a much simpler “n” problem, with each “n” being 1-versus-others. This avoids the combinatorial explosion inherent to the analysis of multi-individual interactions.

A key prerequisite for social role identification is to reliably track every individual without identity switching. As demonstrated throughout the Results, we eliminated identity switching by training the *Detector* module with visually challenging video frames, for example, with overlapping or huddling animals. Importantly, relatively low amounts of manually labeled video frames were sufficient to eliminate identity switching. We recognize that multi-animal tracking remains challenging for computational ethology, and real-world deployments of our *Detector* module may undergo some degree of identity switch, especially when analyzing long-duration videos (e.g., with several hours). However, the fact that we obtained 100% identity consistency from challenging videos is encouraging, and makes us believe that the *Detector* module will work for most applications.

Another known aspect of supervised-learning tools for behavioral classification, such as LabGym2, is the necessity of labeling behaviors for neural network training, which can be labor-intensive. However, LabGym2 has several features that mitigate that labor, such as *Spotlight Animation* and *Pattern Image*, customizable neural networks, and data augmentation. Together, these features increase the efficiency of deep learning, thus reducing the number of behavioral examples needed for training. As demonstrated throughout the Results, only around 200 examples per behavioral category were needed to train an accurate *Categorizer*, which can then be deployed to the rest of the scientific project (or projects). In addition, LabGym2 includes user interfaces that facilitate the manual sorting of behavioral examples, accelerating the construction of the training datasets.

LabGym2 and all of its modules (*Detector, Spotlight Animation* and *Pattern Image*, and *Categorizer*) are subject to continuing and future software development, given their open-source availability (https://github.com/umyelab/LabGym). Future improvements can aim at achieving identity consistency with lower annotation effort. We also expect improvements to the graphical user interface: from network training to analysis outputs. For example, future iterations of LabGym could include the integration of behavior timestamps with other recording modalities, such as brain activity, physiological signals, and vocalizations, enabling the multi-dimensional analysis of complex behaviors.

## METHODS

### Overall implementation

LabGym2 was written in Python programming language (version 3.9)^44^. Python libraries used in LabGym2 are: Numpy^45^, Scipy^46^, scikit-learn^47^, scikit-image^48^, Matplotlib^49^, OpenCV^50^, Pandas^51^, Seaborn^52^, Tensorflow^53^, wxPython^54^, PyTorch^55^, Detectron2^18^.

### Computation hardware

The computational procedures in this study were performed on the following computer systems. (1) University of Michigan Great Lakes Clusters, 2.4 GHz Intel Xeon Gold 6148 with 180 GB RAM, NVIDIA Tesla V100 with 16 GB VRAM. (2) University of Michigan Great Lakes Clusters, 2.9 GHz Intel Xeon Gold 6226R with 372 GB RAM, NVIDIA A40 with 48 GB VRAM. (3) Dell Alienware m16 R1, Windows 11 Home, Intel Core i9 13900HX with 64 GB RAM, NVIDIA GeForce RTX 4090 with 16GB VRAM.

### *Detector* module

#### Post-processing of Detectron2

The video frames are input to Detectron2, which then generates the masks and names of all detected animals or objects in the frames. The name of a mask indicates which animal or object category it belongs to. The names are defined by users when they annotate the images for training a Detector. To filter out overlapping masks that track the same object, the outputs of Detectron2 are filtered by a non-maximum suppression-based method. Briefly, if the intersection area of two masks is greater than 80% of the larger mask, the two masks are considered as overlapping masks tracking the same object. In that case, only the smaller mask is excluded. All masks are then used to generate the foreground of animals or objects in frames. The foreground includes all the pixels inside the contour of the animal or object.

#### Tracking the animals or objects

A trained *Detector* outputs the foreground and the name of a detected animal or object. The foreground includes all the pixels inside the contour of the animal or object and the name indicates which animal or object category this foreground belongs to. Each detected animal or object is assigned a nested python dictionary for storing the frame-wise information, such as name, identity, contours, and centers. The identity is a number that is randomly assigned at the first frame of the video, which is used to distinguish different individuals of the same name. Each object is tracked using an appearance- and distance-based method. Briefly, in two consecutive frames, objects with the same name (similar appearance) are considered as the same animals or objects. If multiple individuals with the same name are present in a frame, the Euclidean distance between every possible pair of two centers in consecutive frames is calculated, and the shortest distance links the two centers that are likely to be the same object in consecutive frames.

### Training of *Detectors*

All *Detectors* were trained using LabGym2’s graphical user interface (GUI).

#### *Detectors* for the mating choice assay with 5 *Drosophila* adults

*Drosophila* adults may have different appearance features such as eye colors that can be used to distinguish individuals. In that scenario, users may use different names such as “red-eye fly” and “white-eye fly” when annotating training images to enhance the distinguishing accuracy and avoid identity switch. However, the resolution of our 5-fly videos was insufficient to distinguish individual flies using their appearance features. Therefore, we defined only one class, “fly”, when annotating the training images. To generate behavior examples (*Spotlight Animations* and *Pattern Images*), we annotated 50∼100 frames out of 15,000 frames for each training video (6 videos in total), and trained a *Detector* for each video with the inferencing frame size of 640 (the original video frame size is 640 × 640) and 5,000 training iterations. These *Detectors* typically had less than 5 identity switches in 10-minute video duration. The behavior examples with identity switches were discarded during behavior-example sorting. During testing, to eliminate identity switching, we annotated 200∼300 frames out of 15,000 frames for each testing video (12 videos in total), and trained a *Detector* for each video with the inferencing frame size of 640 (the original video frame size is 640 × 640) and 5,000 training iterations. The training of a *Detector* typically took less than 10 minutes to complete with any of the three computer systems mentioned above.

#### *Detectors* for the paired prairie voles

The prairie voles in our videos looked identical to each other. Therefore, we defined only one class, “vole”, when annotating the training images. To generate behavior examples (*Spotlight Animations* and *Pattern Images*), we annotated 20∼100 frames out of 18,000 frames for each training video (7 videos in total), and trained a *Detector* for each video with the inferencing frame size of 800 (the original video frame size is 800 × 450) and 5,000 training iterations. These *Detectors* typically had less than 3 identity switches in 10-minute video duration. The behavior examples with identity switches were discarded during behavior example sorting. To eliminate identity switching during the human-LabGym2 benchmarking comparison, we annotated 300∼400 frames out of 18,000 frames for each testing video (2 videos in total), and trained a *Detector* for each video with the inferencing frame size of 800 (the original video frame size is 800 × 450) and 5,000 training iterations. To quantify behavioral differences between the Ctrl and ELSD groups, we trained one *Detector* for all videos (15 videos in total) on all annotated frames. Identity switching was tolerated in these Ctrl vs. ELSD comparisons, as they focused on the quantification of behavioral events irrespective of social roles. The training of a *Detector* typically took less than 10 minutes to complete with any of the three computer systems mentioned above.

#### *Detectors* for interactions between rat and light cue

Each rat was visually distinguishable from the light cue in our videos. Therefore, we defined two classes, “rat” and “light”, when annotating the training images. Since the animals or objects in this dataset had distinct features for the *Detectors* to distinguish, we annotated only 100 frames out of 150,000 frames for 10 videos (5 training videos, 5 testing ones) and trained one *Detector* with the inferencing frame size of 480 (the original video frame size is 480 × 240) and 5,000 training iterations. The *Detector* training took less than 10 minutes to complete with any of the three computer systems mentioned above. This *Detector* achieved 100% detection and tracking accuracy (0 identity switching) in all 10 rat videos in this study.

#### *Detectors* for interactions between mice and objects

Each mouse was visually distinguishable from the two objects in the cage, i.e., the two stimulus mouse holders. In addition, the two objects were distinguishable from each other by the presence vs. absence of a hand-drawn cross. Therefore, we defined three classes: “mouse”, “blank object” and “cross-marked object” when annotating the training images. Since the animals or objects in this dataset had distinct features for the *Detectors* to distinguish, we annotated ∼400 frames out of 1,845,000 frames (123 videos, 7 training videos, 116 testing ones), and trained one *Detector* with the inferencing frame size of 960 (the original video frame size is 960 × 540) and 5,000 training iterations. The *Detector* training took less than 10 minutes to complete with any of the three computer systems mentioned above. This *Detector* achieved 100% detection and tracking accuracy (0 identity switching) in all 123 mouse videos in this study.

#### *Detectors* for macaques in the field

The macaques in our videos looked identical to one another. Therefore, we defined only one class, “macaque”, when annotating the training images. To eliminate identity switching, we annotated 30∼40 frames out of 750 frames for each video (8 videos in total, 5 training videos and 3 testing ones), and trained a *Detector* for each video with the inferencing frame size of 960 (the original video frame size is 960 × 540) and 5,000 training iterations. The training of a *Detector* typically took less than 10 minutes to complete with the three computer systems mentioned above.

### *Spotlight Animations* and *Pattern Images*

The *Spotlight Animation* of a group of interacting animals (or objects) consists of a set of blob-like shapes (including contour outlines) over a user-defined time window encompassing the behavior. Each blob-like shape is a cropped video frame containing all pixels of an individual animal or object. Every individual in the social group is iteratively assigned the “main character” role, whose blob pixels are kept in the RGB color scale of the original video. We then include a magenta-colored (255, 0, 255) outline around the “main character” blob, indicating which animal (or object) is in the *Spotlight* in that iteration. All other blobs (i.e., the “other characters”) have their pixels shown in gray scale. We decided to use magenta for the *Spotlight* outline to maximize contrast, and because the magenta color is rare among animals.

For each *Spotlight Animation* we produce a corresponding *Spotlight Pattern Image*. The *Spotlight Pattern Image* was generated by drawing all the contours of detected animals or objects within the behavior episode onto a black background. The contours of the *Spotlight* individual are then shown in a gradually changing color scale from blue to red, with blue indicating the beginning and red indicating the end of the temporal sequence. All other contours (i.e., the “other characters”) are shown in gray scale.

Therefore, a social group of “n” animals or objects yields “n” iterations of *Spotlight Animations* and *Spotlight Pattern Images*. In each of these iterations, only one individual (“main character”) is assigned *Spotlight* colors. Thus, the same time window or behavioral event looks different depending on which individual is assigned the “main character” color code. This allows us to sort behavioral categories based on the social role of the “main character” and enables the deep neural network to learn and distinguish the social role of each individual. To further enhance the efficiency of the analysis, users can set a “social distance” criterion to exclude animals or objects considered irrelevant to a given social interaction event. If a given animal or object is located beyond the “social distance” criterion (i.e., if it is too distant from the behavioral event and its “main character”), then that animal or object will be excluded from the *Spotlight Animation* and *Pattern Image*.

All the *Spotlight Animations* and *Pattern Images* were generated using LabGym2’s GUI.

### *Categorizer* module

The *Categorizer* module is modified from LabGym1^19^. Briefly, The *Categorizer* contains three submodules: *Animation Analyzer* to analyze *Spotlight Animations, Pattern Recognizer* to analyze *Spotlight Pattern Images*, and *Decision Maker* to determine the behavioral category of each *Spotlight Animation* and *Pattern Image*. All three submodules of the *Categorizer* consist of deep neural networks designed for their analysis purposes. Specifically, the *Animation Analyzer* initially uses time-distributed convolutional layers to analyze raw pixel values representing the animals or objects in each frame of an *Animation*, in order to learn its frame-wise spatial details. The *Animation Analyzer* then uses recurrent layers to compute temporal connectivity among these frame-wise spatial details and learn how they are organized along the temporal axis. The *Pattern Recognizer*, on the other hand, comprises convolutional layers and focuses on the overall motion patterns of the animals or objects during the behavior. Finally, The *Decision Maker* uses a concatenating layer to integrate the outputs from both the *Animation Analyzer* and the *Pattern Recognizer*, and transmits the integrated information to fully-connected (dense) layers for discriminating behavioral categories.

The *Categorizer* additionally includes the “social distance” implementation, which can be set by users to exclude the individuals that are likely not interacting with the *Spotlight* individual (the “main character”). If the distance between an animal (or object) and the “main character” is greater than the “social distance” criterion, that animal or object will be excluded from both the *Spotlight Animation* and *Pattern Image*. The “social distance” criterion is an integer value related to the average body length of all individuals in a video. For example, if the social interaction is deemed unlikely when an individual is distant from the “main character” by more than two-fold the average body length, then users can set the “social distance” value to “2”. If users desire to always include all individuals, they can set this value to “0”, which is decoded by LabGym2 as “infinity”.

The users can flexibly train the *Categorizer* to match the level of complexity of their dataset and behavioral categories. The number of behavioral examples needed to train each *Categorizer* in this study were listed in the figures accordingly. All *Categorizers* used here were trained using LabGym2’s GUI. The training of a *Categorizer* typically took less than 2 hours to complete with the three computer systems mentioned above.

#### *Categorizer* for testing the saliency of *Spotlight* approach

We trained 3 types of *Categorizers* (social distance: 2) with three different complexity levels on the sorted behavior examples (*Pattern Images*, 15 frames as duration at 25 fps) and selected the settings that achieved best training performance (smallest validation loss). The three different complexities were: 1. Pattern Recognizer (LV2, 32 × 32 ×3). 2. Pattern Recognizer (LV3, 64 × 64 ×3). 3. Pattern Recognizer (LV4, 64 × 64 ×3). No.3 was selected, and was trained for 3 repeats. The repeat with smallest validation loss was used for analysis.

#### *Categorizer* for courtship choice in 5 *Drosophila* adults

We trained 3 types of *Categorizers* (social distance: 2) with three different complexity levels on the sorted behavior examples (*Spotlight Animations* and *Pattern Images*, 15 frames as duration at 25 fps) and selected the settings that achieved best training performance (smallest validation loss). The three different complexities were: 1. Animation Analyzer (LV2, 32 × 32 × 3) + Pattern Recognizer (LV3, 64 × 64 ×3). 2. Animation Analyzer (LV5, 64 × 64 × 3) + Pattern Recognizer (LV5, 64 × 64 ×3). 3. Animation Analyzer (LV5, 64 × 64 × 3) + Pattern Recognizer (LV5, 96 × 96 ×3). No.2 was selected, and was trained for 3 repeats. The repeat with smallest validation loss was used for analysis.

#### *Categorizer* for dyadic interactions in prairie voles

We trained 3 types of *Categorizers* (social distance: 2) with three different complexity levels on the sorted behavior examples (*Spotlight Pattern Images*, 20 frames as duration at 30 fps) and selected the settings that achieved best training performance (smallest validation loss). The three different complexities were: 1. Pattern Recognizer (LV2, 64 × 64 ×3). 2. Pattern Recognizer (LV4, 64 × 64 ×3). 3. Pattern Recognizer (LV4, 96 × 96 ×3). No.3 was selected, and was trained for 3 repeats. The repeat with smallest validation loss was used for analysis.

#### *Categorizer* for social recognition in mice

We trained 3 types of *Categorizers* (social distance: 2) with three different complexity levels on the sorted behavior examples (*Spotlight Animations* and *Pattern Images*, 15 frames as duration at 15 fps) and selected the settings that achieved best training performance (smallest validation loss). The three different complexities were: 1. Animation Analyzer (LV2, 64 × 64 × 3) + Pattern Recognizer (LV3, 64 × 64 ×3). 2. Animation Analyzer (LV3, 64 × 64 × 3) + Pattern Recognizer (LV4, 64 × 64 ×3). 3. Animation Analyzer (LV4, 64 × 64 × 3) + Pattern Recognizer (LV4, 128 × 128 ×3). No.3 was selected, and was trained for 3 repeats. The repeat with smallest validation loss was used for analysis.

#### *Categorizer* for social interactions in wild macaques

We trained 3 types of *Categorizers* (social distance: 2) with three different complexity levels on the sorted behavior examples (*Spotlight Animations* and *Pattern Images*, 25 frames as duration at 25 fps) and selected the settings that achieved best training performance (smallest validation loss). The three different complexities were: 1. Animation Analyzer (LV2, 32 × 32 × 3) + Pattern Recognizer (LV3, 32 × 32 ×3). 2. Animation Analyzer (LV2, 32 × 32 × 3) + Pattern Recognizer (LV3, 64 × 64 ×3). 3. Animation Analyzer (LV3, 64 × 64 × 3) + Pattern Recognizer (LV3, 64 × 64 ×3). No.2 was selected, and was trained for 3 repeats. The repeat with smallest validation loss was used for analysis.

### Criteria for sorting the behavior examples

#### Drosophila

*Orienting*: A fly is repositioning its body toward a conspecific.

*Singing*: A fly is extending and vibrating one of its wings.

*Attempted copulation*: A fly is bending its abdomen and trying to copulate with a conspecific.

*Being courted*: A fly is being oriented, singed or receiving copulation attempts from a conspecific.

*Idling*: A fly is immobile and is not interacting with conspecifics.

*moving*: A fly is moving and is not interacting with conspecifics.

#### Prairie vole

*Approaching*: A prairie vole is moving toward the conspecific that is idling in place.

*Sniffing*: A prairie vole is sniffing or closely investigating the conspecific.

*Being sniffed*: A prairie vole is being sniffed or closely investigated by the conspecific.

*Huddling*: The two prairie voles are immobile or showing small in-place movements while maintaining side-by-side body contact regardless of body direction.

*Aggression*: A prairie vole is either: (1) extending its anterior paws or violently jumping toward the conspecific; or (2) escaping from such aggressive actions.

*Moving*: A prairie vole is moving alone without interacting with the conspecific.

*Idling*: A prairie vole is immobile or showing small in-place movements without interacting with the conspecific.

*Chasing*: A prairie vole is moving toward the conspecific that is also moving in the same direction, forming overlapping trajectories.

*Being chased*: A prairie vole is escaping from the conspecific that is also moving in the same direction, forming overlapping trajectories.

A simplified behavioral category system contained two categories: *Combined social*, and *Combined non-social*. The *Combined social* category included *Approaching, Sniffing, Being sniffed, Huddling, Aggression, Chasing*, and *Being chased*. The *Combined non-social* category included *Moving* and *Idling*.

#### Mouse

*Climbing on blank*: The mouse is climbing the blank object.

*Climbing on cross-marked*: The mouse is climbing the cross-marked object.

*Rearing on blank*: The mouse is rearing on the blank object.

*Rearing on cross-marked*: The mouse is rearing on the cross-marked object.

*Sniffing blank*: The mouse is sniffing the blank object.

*Sniffing cross-marked*: The mouse is sniffing the cross-marked object.

*Moving*: The mouse is roaming distant from either object.

*Resting*: The mouse is immobile or showing small in-place movements distant from either object.

*Rearing*: The mouse is rearing on the walls of the arena.

A simplified behavioral category system contained three categories: *Combined cross-marked, Combined blank*, and *Combined away*. The *Combined cross-marked* category included *Climbing on cross-marked, Rearing on cross-marked*, and *Sniffing cross-marked*. The *Combined blank* category included *Climbing on blank, Rearing on blank*, and *Sniffing blank*. The *Combined away* category included *Moving, Resting*, and *Rearing*.

#### Macaque

*Grooming*: A macaque is using its hands to groom a conspecific.

*Being groomed*: A macaque is being groomed, either sitting or lying on the ground, by one or more conspecifics.

*Others*: A macaque is performing a behavior other than grooming or being groomed, such as moving or foraging.

### Behavioral experiments and recordings

#### Courtship choice in *Drosophila*

We used a subset of videos of *Drosophila* mating choice assay from a previous report^23^. In these experiments, a male *Drosophila* adult was paired with 4 females, 2 younger and 2 older, in a lab-made mating chamber (1.5 cm in diameter and 0.3 cm deep) and recorded for 10 minutes at 25 frames per second. The male *Drosophila* adults could either express irrelevant control proteins (Ctrl group) or neurodegenerative proteins (Exp group) in their central nervous system. All videos were resized to 640 × 640 pixels using a preprocessing module in LabGym2’s GUI.

#### Pavlovian Cocaine Cue Task

Animal housing and ethics were approved by the institution involved in the rat experiments (University of Michigan LSA Psychology: IACUC PRO00011037). Twenty Sprague Dawley rats, evenly distributed across sexes and weighing 250–275 grams, were individually housed in cages. They were maintained on a 12-hour light/dark cycle with controlled temperature and humidity upon arrival (procured from Envigo and Taconic). The rats had unrestricted access to Laboratory Rodent Diet 5001 (LabDiet) and water. A one-week acclimation period in the colony room preceded any experimentation, during which the rats were regularly handled. All subsequent training sessions were conducted during the 12-hour lights-on period. Approval for all procedures was obtained from the University of Michigan Institutional Animal Care and Use Committee, and the experiments were carried out in laboratories accredited by the Association for Assessment and Accreditation of Laboratory Animal Care. The rats were subjected to tests in conditioning chambers (20.5 × 24.1 cm floor area, 20.2 cm high; MED Associates Inc.) featuring a cue light (13.5 cm above floor) on the wall opposite to the red house light. Additionally, the chambers were equipped with overhead video cameras (frame size: 1024 × 768; video format: .avi), and behavior was recorded throughout each session. All videos were resized to 480 × 240 pixels using a preprocessing module in LabGym2’s GUI.

#### Dyadic interactions in prairie voles

Details on animals and video recordings have been published by a subset of the present coauthors^26,27^. Briefly, animal housing and ethics were approved by the two institutions involved in the prairie vole experiments (University of Michigan Medical School: IACUC PRO00009818; Veterans Affairs Portland Health Care System: ACORP 3652-18). For induction of early-life sleep disruption (ELSD)^26^, bedded home-cages containing prairie vole pups (14-21 days of age) and their parents were gently and intermittently agitated using an orbital shaker (agitated for 10 seconds every 110 seconds, 110 rotations per minute). Pups were then weaned into groups of 2-4 same-sex siblings and co-housed until adulthood (>20 weeks of age), when recordings occurred. Control animals (Ctrl) were exposed to the same housing conditions but without orbital shaking.

Sexually naïve male-female pairs were randomly formed from these adult animals into all possible sex vs. group combinations (Male-Ctrl/ Female-Ctrl, Male-ELSD/Female-ELSD, Male-Ctrl/Female-ELSD, Male-ELSD/Female-Ctrl). Each male-female pair was placed in a bedded home-cage with a wire-mesh divider at the center, confining each individual to its own non-overlapping space. The animals were then filmed (camera model: Basler, acA1300-60gm, infrared illumination) from an overhead angle (90°) for 72 h (14 h/10 h light/dark cycle; lights on at 5:00 am; cameras on at 8:00 am) while they interacted through the divider, producing spatial and temporal variables of pair bonding behavior, analyzed in another study^27^.

The present study used separate videos from the same animals, recorded after pair bonding (4 days after the 72 h recording), with shorter duration (3-4 h) and without the divider. This allowed the animals to engage in the freely-moving dyadic interactions we examined with LabGym2. Because the focus of the present study is on software development, we simplified our groupings (only Male-Ctrl/Female-Ctrl and Male-ELSD/ Female-ELSD pairs were used) and statistical comparisons (Ctrl vs. ELSD, males and females combined). Furthermore, only the initial 10 minutes of each video were analyzed, just enough to capture the transition from high behavioral activity (initial ∼5 minutes period after placing the animals in the cage) to quieter states with higher incidence of huddling. All videos were cropped to the frame size of 800 × 450 pixels and contrast-enhanced using a preprocessing module in LabGym2’s GUI.

#### Social recognition in mice fed with high-fat diet

Animal housing and ethics were approved by the Feldman protocol, PRO00010039. Sixty male C57BL/6J mice (The Jackson Laboratory) were obtained, 30 of which with 5 weeks of age, and another 30 with 56 weeks of age at time of purchase. Upon arrival, 15 mice from each age group were randomly selected to receive either standard (Ctrl) or high-fat diet (HFD; 60% calories from fat; Research Diets, D12492). Mice were then housed in same-diet and same-age groups of 2-5 individuals in standard conditions (ad libitum food and water, 14 h/10 h light/dark cycle) until the recordings, 24 weeks later. Ctrl and HFD mice were “test mice” to be examined for freely-moving behaviors using LabGym2. An additional cohort of “stimulus mice” was formed with 16 age-matched individuals, housed under the same conditions and fed with standard diet.

The social recognition^56,57^ recordings took place within a rectangular arena (53 × 38 × 26 cm) with two mouse-holding chambers (10 × 10 × 10.5 cm) in opposite corners of the arena. One day prior to the recordings, the test mice were habituated to the arena in two 5-minute periods, 1 h apart. During these periods, the test mice were allowed to explore the arena and two empty chambers from the outside, i.e., without being able to enter these chambers. Stimulus mice, in turn, were habituated by being held inside one of the chambers for four 5-minute periods, 1 h apart. One day after habituation, the animals were filmed (camera model: Logitech, C615 HD Webcam, white light illumination) from an overhead angle (90°) during two 10-minute sessions: session #1 and session #2. In session #1, a stimulus mouse was placed in one of the holding chambers and the test mouse was allowed to explore. That was followed by a memory retention interval (5 minutes) when the test mouse was removed from the arena. The test mouse was then put back into the arena for session #2, which included a novel mouse in the previously empty chamber. Test and stimulus mice were able to exchange olfactory cues through fine mesh cutouts around the holding containers. All videos were resized to 960 × 540 pixels and contrast-enhanced using a preprocessing module in LabGym2’s GUI.

#### Social interactions in wild Japanese macaques

The macaque videos were recorded by Cédric Sueur in January 2024 at two sites: the Arashiyama Monkey Park in Kyoto, Japan^58,59^ and the Choshikei Monkey Park on Shodoshima, Japan^60,61^. These videos feature Japanese macaques, which are wild but provisioned. These groups have become accustomed to park staff and visitors, allowing for observations from less than 5 meters away without disturbing them. Although their diet includes natural forest foods, it primarily consists of food provided by the park: wheat grains distributed three times a day (approximately 20 kg per group in total) and sweet potatoes and vegetables offered in the afternoon. Tourists also occasionally feed the monkeys small amounts of peanuts and soybeans. All adult monkeys are recognized and identified, but the juveniles and subadults are not.

Grooming behaviors^41,42^ were recorded using behavioral sampling during 30-second sessions, with grooming pairs chosen at random. The equipment used included a Canon EOS 5D Mark IV camera with a 105-mm Canon lens. This footage contributes to two studies aiming to automatically identify individuals and behaviors using deep learning to build social networks^62,63^.

### Testing *Categorizers* with unseen videos

LabGym2 has a functional unit called “Test *Categorizers*”, which calculates performance metrics (precision, recall, specificity, f1-score, and overall accuracy) of a trained *Categorizer* on the ground-truth testing data. To establish a ground-truth testing dataset, the behavior examples (*Spotlight Animations* and *Pattern Images*) were generated from the testing videos (different from the training videos) using LabGym2’s GUI. The behavior examples were then sorted by human experts. The sorted behavior examples (ground-truth testing dataset) were input into the Test *Categorizers* functional unit for testing.

The performance metrics were calculated as below: precision (equation 1), recall (equation 2), specificity (equation 3), f1 score (equation 4).

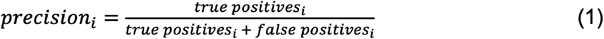

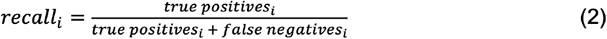

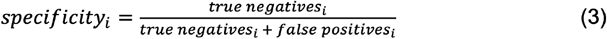

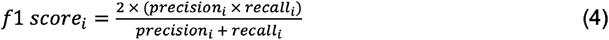

### Quantification and statistics

All quantitative measures of each behavior were automatically calculated by LabGym1’s *Quantifier* Module, as reported in detail previously^19^. These quantitative measures are taken as summary values from the entire analyzed period (e.g., 10 minutes) and are calculated as follows:

1. *Count* is the number of occurrences of a behavior. Consecutive occurrences of the same behavior are counted as one.
2. *Distance* is the total distance traveled by the animal while performing a given behavior.
3. *Duration* is the total time performing a given behavior.
4. *Intensity (area)* measures changes in animal body area (*a*) between frames divided by the time window (*t*) during which a behavior was performed (equations 5).

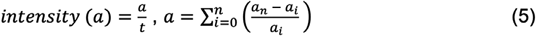
5. *Speed* measures the distance (*d*) traveled by the animal (its center of mass) between adjacent frames divided by the time window (*t*) during which a behavior was performed (equation 6).

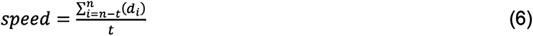
6. *Velocity* measures the shortest distance between the beginning and end location (*dt*) of the animal (its center of mass) divided by the time (*t*_*occuring*_) during which such displacement occurs (equation 7).

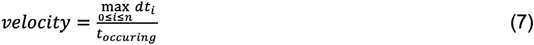
7. *Vigor (area)* measures the magnitude (area) divided by the time (*t*_*occuring*_) during which that change occurs (equations 8). *Magnitude (area)* measures the maximum proportional change in animal body area (*a*) when performing a behavior.

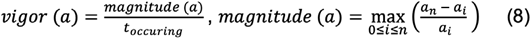

Statistical tests were also performed using LabGym2’s GUI. Briefly, after the “control” and “experimental” groups were selected through the GUI, LabGym2 automatically examined the distribution of the data using Shapiro–Wilk normality test and then performed either parametric or non-parametric tests accordingly. These tests are indicated in their corresponding figure legends.

## ACKNOWLEDGEMENTS

We thank Drs. Linus Manubens-Gil for helpful discussions. This research was supported by National Institutes of Health (NIH) to B.Y. (R01NS128500, R01NS104299), B.O.W.-M.M.L. (R01MH131592), C.R.F. (R01DK13024, R01DA044204), and Portland VA Research Foundation to B.O.W.- M.M.L.. The content is solely the responsibility of the authors and does not necessarily represent the official views of the NIH or Department of Veterans Affairs of the United States Government.

## AUTHOR CONTRIBUTIONS

B.Y. and Y.H. conceived the project. B.Y. supervised the project. Y.H. designed and programmed LabGym2 with the assistance of K.G., R.S., and I.B.. L.S.B.J., K.S., G.G.M., and H.C. performed the experiments in rodents. K.G., L.S.B.J., K.S., T.A., H.C., and Y.H. performed data labeling and analysis. C.E.J.T. and M.M.L. provided rodents. C.S. recorded macaque videos. L.S.B.J., K.S., J.Z., M.M.L., B.O.W., C.S., C.R.F., B.Y., and Y.H. wrote the manuscript.

## DECLARATION OF INTERESTS

The authors declare no competing interests.

## DATA AND CODE AVAILABILITY

The datasets used in this study will be deposited to Zenodo and be publicly available. The code generated in this study is publicly available in GitHub: https://github.com/umyelab/LabGym. The first version of LabGym2 (LabGym2.0) was released via GitHub in July, 2023.

## EXTENDED DATA

**Extended Data Figure 1.**
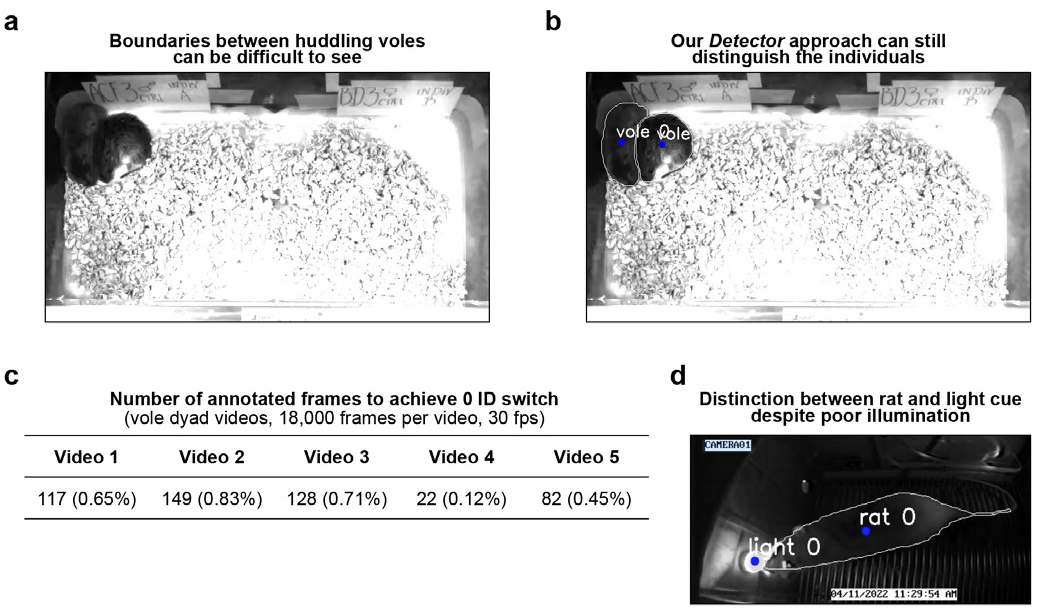
LabGym2’s Detectors requires low amount of training frames and can accurately track individuals in various challenging scenarios. **a**, An example snapshot showing that the boundaries between huddling voles can be difficult to see. **b**, An example snapshot showing that a *Detector* trained on 22 annotated frames can still segment the two huddling voles very well. **c**, Each vole dyad video required a relatively low amount of annotated frames to achieve 0 identity switch. **d**, An example snapshot showing that a *Detector* trained on 100 annotated frames can generalize to unseen videos and precisely segment a rat versus a light cue, despite low illumination of the experimental chamber and poor-resolution of the video.

## Notes

https://github.com/umyelab/LabGym

### Competing Interest Statement

The authors have declared no competing interest.

